# Toxicity of 2 pg ethynylestradiol in brown trout embryos (*Salmo trutta*)

**DOI:** 10.1101/161570

**Authors:** Lucas Marques da Cunha, Anshu Uppal, Emily Seddon, David Nusbaumer, Etienne L. M. Vermeirssen, Claus Wedekind

## Abstract

Endocrine disrupting chemicals are a threat to natural fish populations in the aquatic environment. Their toxicity is usually discussed relative to concentrations in the water the fish are exposed to. In the case of the synthetic compound 17-alpha-ethynylestradiol (EE2), a common and persistent estrogen, concentrations around 1 ng/L have repeatedly been found to induce toxic effects in fish. Here, we used brown trout (*Salmo trutta*) from a natural population to study EE2 take up and how it affects early life-history. We collected adults during the spawning season, produced 730 families *in vitro* (to control for potential maternal and paternal effects on embryo stress tolerance), and singly raised 7,300 embryos (in a 2 mL static system) that were either exposed to one dose of EE2 at 1 ng/L (i.e., 2 pg/embryo) or sham-treated. We found that EE2 concentration did not significantly change over a period of 3 months in control containers without embryos. Embryos took up most of the 2 pg EE2 within about 4 weeks at 4.6°C. EE2 treated embryos experienced higher mortality, delayed hatching of the survivors, and had reduced size at hatching. Our findings suggest that the toxicity of EE2 is often underestimated when discussed at the level of concentrations in water only.

## Introduction

Pollution by endocrine disrupting chemicals is a threat to wildlife (Wedekind, 2017). The synthetic estrogen 17α-ethynylestradiol (EE2) is a persistent and powerful endocrine disruptor (reviewed in Corcoran et al., 2010, Aris et al., 2014). This steroid is a compound of oral contraceptive pills and reaches the environment through household sewage (Ternes et al., 1999a, Chèvre, 2014). During the sewage treatment process, EE2 removal is expected to be on average 68%, i.e. its degradation is likely not complete (Johnson et al., 2013). Thus, rivers that carry treated sewage effluent also carry EE2. Modelling work suggests that EE2 is present at 10 pg/L or higher in 20% of the entire European river network (Johnson et al., 2013) and concentrations around 1 ng/L have been regularly measured in surface waters (Tiedeken et al., 2017).

In fish, EE2 exposure is believed to have two main types of effects. First, it may affect early life-history stages, when individuals are often more sensitive than at later stages, by altering traits such as embryo survival, developmental time and larval growth (reviewed in Aris et al., 2014). Secondly, as EE2 is a hormone and sex differentiation in fish can often be affected by environmental factors (Devlin & Nagahama, 2002), EE2 can disturb fish reproductive traits in juveniles and mature individuals (reviewed in Leet et al., 2011, Caldwell et al., 2012). Either type of effect has potential consequences for individual and population fitness (Cotton & Wedekind, 2009, Wedekind, 2017). Significant effects have been observed in concentrations around 1 ng/L (Caldwell et al., 2012, Aris et al., 2014). For example, in zebrafish (*Danio rerio*) an exposure to 1 ng/L EE2 increased vitellogenin body content (Van den Belt et al., 2003) and 2 ng/L were sufficient to reduce embryo survival (Santos et al., 2014) and growth (Xu et al., 2008). Moreover, juvenile medaka (*Oryzias latipes*) exposed to 10 ng/L of EE2 showed reduced survival and growth (Scholz & Gutzeit, 2000). Similar concentrations have been linked to the appearance of gonadal deformations in mature zebrafish males (Xu et al., 2008) and led to intersex medaka males (Metcalfe et al., 2001). The toxicity of EE2 has, however, rarely been studied in non-model fish. While model species offer good background for ecotoxicology tests, understanding the toxicity of compounds with wild non-model species is an important step in validating results in a more ecologically-relevant context (Wedekind et al., 2007). Moreover, fish ecotoxicology studies often focus only on the effects of constant concentrations of chemical compounds in the water (OECD, 1992, Leet et al., 2011, Aris et al., 2014), but see Bjerregaard et al. (2008) and Knudsen et al. (2011). However, determining both, exposure content and individual uptake, is necessary to better understand the toxicity of a substance (Quinnell et al., 2004, Skillman et al., 2006).

Species from the salmonidae family are typically of high economic, cultural and ecological relevance. Nonetheless, salmonids have been suffering from anthropogenic changes in the environment and several populations face demographic shifts (e.g. Wedekind et al., 2013) and population declines (e.g. Gustafson et al., 2007, Wedekind & Küng, 2010). Because many salmonid species spawn in rivers – many of which affected by effluent discharges – and embryos typically develop in an immobile stage over a period of several months, EE2 may be a particular threat for early life stages of these species. Moreover, EE2 is likely a stronger threat in rivers than in lakes because it typically reaches higher concentrations in smaller rivers that receive high effluent loads. When Brazzola et al. (2014) exposed two species of whitefish (*Coregonus palaea* and *C. albellus*) to low concentrations of EE2 (from 1 ng/L), they found EE2 to reduce embryo survival and larval growth and to delay hatching (Brazzola et al., 2014). However, whitefish are lake-spawning salmonids that in the natural environment may be less exposed to EE2 than their river-spawning counterparts. Schubert et al. (2014) investigated the effects of exposures to the natural estrogen 17β-estradiol (E2) on embryos of brown trout (*Salmo trutta*), a river-spawning salmonid. The authors found an E2 exposure of 3.8 and 38 ng/L to not significantly reduce embryo survival, but to delay hatching time and reduce body length at hatching (Schubert et al., 2014). However, the trout population used in Schubert et al. (2014) was captive. Because salmonids display large maternal effects (Einum & Fleming, 1999, Clark et al., 2014) that include environmentally-dependent compounds that females allocate to their eggs (e.g. Wilkins et al., 2017, Wilkins et al. submitted manuscript), the toxicity of chemicals in an ecologically-relevant context requires investigations on wild individuals.

The brown trout population of the Aare river system in Switzerland has been extensively monitored and has shown a decline of about 50% over the past three decades (Burkhardt-Holm, 2007, Stelkens et al., 2012). The observed population decline is not yet sufficiently understood but habitat fragmentation (Stelkens et al., 2012) and estrogenic pollution (Burkhardt-Holm et al., 2008) have been investigated as candidates contributing to it. In this study, we used brown trout from the Aare river system to investigate (i) whether and how much embryos take up EE2 from a low and ecologically-relevant exposure and (ii) the effects of such uptake on early life-history traits.

## Methods

### Experimental design and embryo rearing

Adult males (N= 142) and females (N=145) were sampled during their spawning season from the Aare river system, in Switzerland. Their gametes were stripped and used for *in vitro* block wise full-factorial fertilizations (Lynch & Walsh, 1998) for a study on variance components and genetic correlations (Marques da Cunha et al., in preparation). Here, the block-wise breeding of gametes of many males and females controls for potential parental effects on stress tolerance. The experimental crosses were performed in 4 different days, once per week from mid-November to mid-December. The breeding block design varied from 4 ×5 (i.e., 4 females crossed with 5 males in all possible combinations to produce 4 ×5 sibgroups) to 6 ×5, depending on the availability of individuals. In total, 29 breeding blocks yielding 730 sibgroups were produced. After hardening for 2 hours, samples of freshly fertilized eggs of each sibgroup were brought to the laboratory where 15 freshly fertilized eggs per sibgroup were used for another study (Marques da Cunha et al., in preparation) and 10 per sibgroup (N = 7,300) were used for the present study. They were washed as in von Siebenthal et al. (2009) and singly raised in polystyrene 24-well plates (Greiner Bio-One, Austria) filled with 1.8 mL of autoclaved standardized water per well (OECD, 1992). Plates were incubated in a climate chamber at 4.6°C.

### Treatment preparation, exposure and trait measurements

A spike solution of 10 ng/L of analytical 17α-ethynylestradiol (Sigma-Aldrich, USA) was prepared through a 3-steps serial dilution. Because EE2 is poorly soluble in water, absolute ethanol (VWR International, USA) was used for the first step of the dilutions. This led to a concentration of 0.004% of ethanol in the EE2 spike solution. Analogously, a control spike solution with the same concentration of ethanol but without EE2 was prepared. All of the dilutions were prepared with autoclaved standardized water (OECD, 1992). One day post fertilization, either 0.2 mL of the EE2 or 0.2 mL of the control spike solution was added to the wells for a final volume of 2 mL. The nominal concentrations in the wells were 1 ng/L of EE2 and 0.0004% of ethanol for EE2-exposed embryos (i.e. a total content per well of 2 pg EE2) or 0.0004% of ethanol for sham-treated individuals.

At the day of hatching, embryos were singly transferred to 12-well plates (Greiner Bio-One, Austria) filled with 3 mL of autoclaved standardized water (OECD, 1992), i.e. there was no EE2 treatment at that stage. These 12-well plates were scanned for embryo body measurements (Epson Perfection V37, Japan), i.e. hatchling length and yolk sac length and width at hatching. After 24 days, the plates were again scanned for the same trait measurements. Larval growth was calculated as larval length at 24 days post hatching minus length at hatching. Yolk sac volume at hatching was calculated as in Jensen et al. (2008). All the trait measurements were performed with ImageJ (http://rsb.info.nih.gov/ij/).

### Statistical analyses

Embryo survival was analyzed as a binomial response variable in generalized linear mixed models (GLMM) and hatching time, hatchling length, yolk sac volume at hatching and larval growth as continuous response variables in liner mixed models (LMM). Treatment was entered as fixed effect and breeding block (which comprises the variation of family and fertilization date) as random effect in the models. The significance of the model terms was obtained by comparing a model including or lacking the term of interest to a reference model with likelihood ratio tests (LRT) and Akaike information criterion (AIC). All the statistical analyses were performed in R (R Development Core Team, 2015) and mixed models with the lme4 package (Bates et al., 2015).

### Determining EE2 concentrations in 24-well plates

A second experiment was performed to estimate embryo EE2 uptake and determine the persistence of EE2 in the same model of polystyrene 24-well plates as used in the first experiment. In total 4,080 brown trout embryos (from other parents of the same populations as in the first experiment; seven 4 ×6 breeding blocks, i.e. 168 sibgroups of 28 females and 42 males) were raised in 170 24-well plates (Greiner Bio-One, Austria) at the same conditions as in the first experiment (i.e. same methods, temperature, treatment and concentrations). Furthermore, 170 24- well plates without embryos were prepared and analogously treated with EE2 or control spike solutions. Measuring EE2 in plates without embryos and comparing these measurements with plates that contanined embryos allowed for estimations of embryo EE2 uptake. Water samples were collected at 5 time points across embryonic development and stored at −20° C for later EE2 measurements. For each time point, the water of 12 entire plates was pooled per treatment (576 mL of water sample per treatment). The first time point was performed only in empty plates and was collected 30 minutes after the spike. The following time points were performed in plates with and without embryos and were collected 7, 28, 56 and 84 days after exposures (the latter was collected a few days before embryo hatching started).

EE2 was quantified with liquid chromatography-tandem mass spectrometry (LC-MS/MS).First, the water samples were thawed and filtered with glass fibre filters. Then, sample volume and pH were set to 250 mL and 3, respectively. After that, 4 ng/L of 17α-ethynylestradiol D4 were added to control for recovery and matrix effects. Water samples were enriched on LiChrolut EN / LiChrolut RP-C18 cartridges (previously conditioned with hexane, acetone, methanol and water at a pH of 3 as in Escher et al. (2008)). Cartridges were dried with nitrogen and eluted with acetone and methanol. Solvents were exchanged to hexane/acetone at a ratio of 65:35 and the extracts were passed through Chromabond Silica columns (Ternes et al., 1999b). Finally, the volume of the extracts was set to 0.25 mL. LC-MS/MS was performed with an Agilent6495 Triple Quadrupole. The column used was an XBridge BEH C18 CP (2.5 μm, 2.1 mm ×75 mm). A gradient of acetonitrile/water was used for the liquid chromatography followed with a postcolumn addition of ammonium fluoride solution. Mass transitions that were monitored are listed in the SI (Table S1). The LC-MS/MS method also covered estrone (E1), 17β-estradiol (E2) and bisphenol A (BPA) – mass transitions described in Table S1 All three compounds were detected in the 24 well-plates with BPA at significant concentrations (see SI).

## Results

### EE2 water quantifications and embryo uptake

The measured concentration of the EE2 and control spike solutions corresponded to the nominal concentrations. The EE2 spike solution concentration was 10.1 ng/L (nominal concentration = 10 ng/L) and the control spike solution concentration was lower than the limit of quantification (LOQ = 0.05 ng/L). The concentrations of EE2 in water samples from 24-well plates depended on the presence of embryos. In EE2-spiked plates without embryos, the concentrations of EE2 remained near the expected nominal concentration (1 ng/L, i.e. a total content of around 2 pg per well) throughout the observational period (Fig. 1). In EE2-spiked plates with embryos, the level of EE2 gradually declined from near the nominal concentration to the limit of quantification (Fig. 1). Nearly all of this decline happened during the first month of embryogenesis (Fig. 1). Control-spiked plates with or without embryos did not show EE2 concentrations above LOQ (which was now at 0.1 ng/L EE2 for these measurements).

**Fig. 1:**
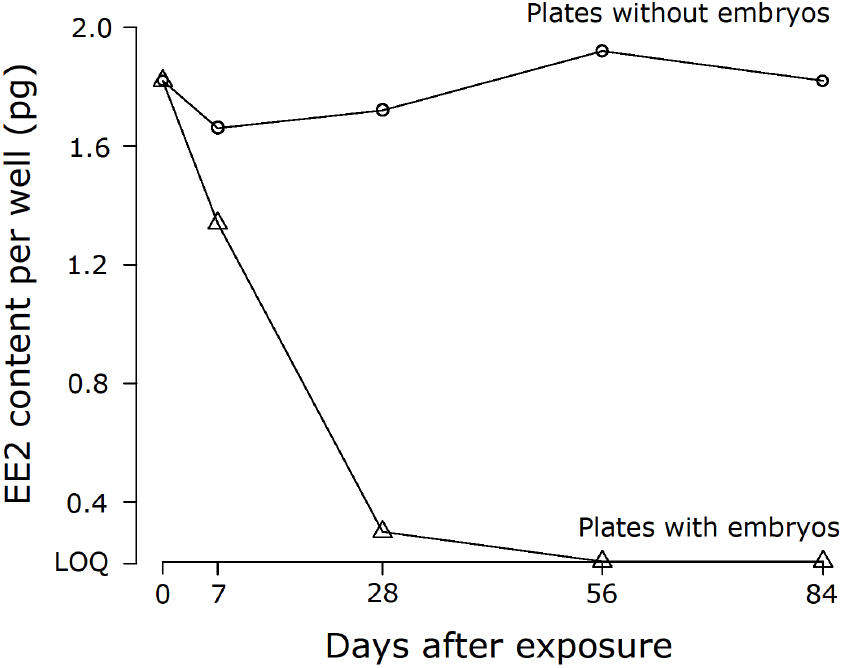
The persistence of 17α-ethynylestradiol (EE2) in 24-well plates with and without embryos across 5 time points. Triangles represent plates with embryos and circles without embryos. Sham-treatment data points are not shown because they were always below LOQ (0.1 ng/L or a total well content of 0.2 pg).

### EE2 effects on brown trout early traits

Exposure to EE2 reduced embryo survival by 0.7%, i.e. an increase in mortality by 34.9 % (Table 1a; Fig. 2a). Breeding blocks varied by their survival rates, however the differences among breeding blocks were not dependent on the treatment (block x treatment interactions in Table 1a). The exposure to EE2 led to delayed hatching time by about half a day (Table 1b, Fig. 2b) and to reduced hatchling length by on average 0.25% (Table 1c; Fig. 2c). Again, breeding blocks differed in their overall hatching time and hatchling length and these effects were not treatment specific (Tables 1b – c). Yolk sac volume at hatching and larval growth were not significantly affected by the EE2 treatment (Tables 1d – e; Fig. 2d – e). Variation in these traits was again strongly dependent on breeding blocks (Table 1d – e).

**Table 1.** The effects of treatment (exposure to EE2) and breeding block on (A) embryo survival,(B) hatching time, (C) hatchling length, (D) yolk sac volume at hatching, and (E) larval growth. Likelihood ratio tests on mixed model regressions were used to compare a reference model (in bold) with models including or lacking the term of interest. Significant effects (p <0.05) are highlighted in bold.

| Model terms | Effect tested | AIC | d.f. | $X^2$ | $P$ |
| --- | --- | --- | --- | --- | --- |
| *(A) Embryo survival* |  |  |  |  |  |
| **t + block** |  | 1219 | 4 |  |  |
| block | t | 1222 | 3 | 4.9 | **0.03** |
| t | block | 1360 | 3 | 142.9 | **<0.001** |
| t + block + t x block | t x block | 1223 | 6 | 0.2 | 0.89 |
| *(B) Hatching time* |  |  |  |  |  |
| **t + block** |  | 29068 | 5 |  |  |
| block | t | 29101 | 4 | 34.8 | **<0.001** |
| t | block | 38808 | 4 | 9742.0 | **<0.001** |
| t + block + t x block | t x block | 29069 | 7 | 3.2 | 0.20 |
| *(C) Hatchling length* |  |  |  |  |  |
| **t + block** |  | 5485 | 5 |  |  |
| block | t | 5488 | 4 | 4.1 | **0.04** |
| t | block | 8113 | 4 | 2629.0 | **<0.001** |
| t + block + t x block | t x block | 5490 | 7 | 0.0 | 0.99 |
| *(D) Yolk sac volume at hatching* |  |  |  |  |  |
| **t + block** |  | 35225 | 5 |  |  |
| block | t | 35226 | 4 | 2.7 | 0.10 |
| t | block | 38521 | 4 | 3297.2 | **<0.001** |
| t + block + t x block | t x block | 35229 | 7 | 0.8 | 0.66 |
| *(E) Larval growth* |  |  |  |  |  |
| **t + block** |  | 5204 | 5 |  |  |
| block | t | 5204 | 4 | 1.3 | 0.26 |
| t | block | 5724 | 4 | 522.1 | **<0.001** |
| t + block + t x block | t x block | 5208 | 7 | 0.5 | 0.77 |

Fixed effect: t, treatment. Random effect: breeding block.

**Fig. 2:**
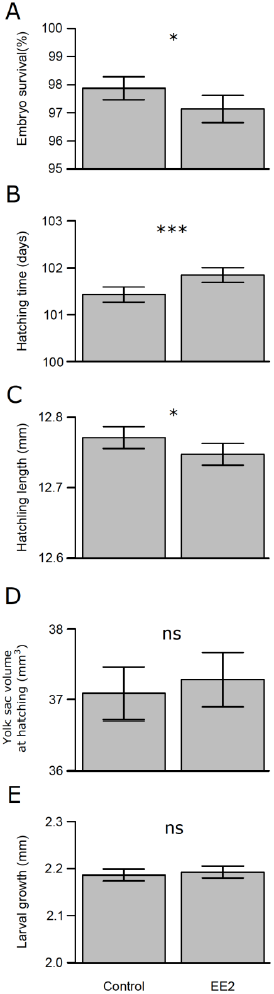
The effects of 17α-ethynylestradiol (EE2) on embryo early phenotype: (A) embryo survival, (B) hatching time, (C) hatchling length, (D) yolk sac volume at hatching, and (E) larval growth. Bars represent means of family means and error bars are 95 % confidence intervals, ^***^= p < 0.001, ^*^ = p <0.05, and ns = p >; 0.05. See Table 1 for statistics.

## Discussion

EE2 is a common and potent endocrine disrupting pollutant of surface waters (Ternes et al., 1999a, Chèvre, 2014). However, its toxicology has mainly been studied on model fish that were bred and raised under laboratory conditions. Studies on wild populations are rare but necessary in this context, because of the many differences in stress tolerance within and among species and between wild and captive populations (Wedekind et al., 2007, Nyman et al., 2014). For example, maternal environmental effects on egg content and hence on embryo viability and stress tolerance typically differ considerably between populations (Pompini et al., 2013, Sopinka et al., 2016, Wilkins et al., submitted manuscript). Moreover, the toxicology of EE2 has mainly been studied relative to concentrations that were typically kept constant in experimental setups, and little is known about the uptake of EE2 in embryos especially of wild populations. We therefore sampled gametes from wild-caught spawners to produce embryos that represent their counterparts in the wild. We exposed them to 2 pg EE2, estimated EE2 uptake and determined various measures of embryo viability in response to this pollutant.

EE2 concentrations in plates without embryos remained constant throughout the observational period of three months. This confirms the high stability and low degradation of EE2 as compared to natural estrogens (Aris et al., 2014). When Ternes et al. (1999a) investigated the behavior and occurrence of estrogens in activated sludge from sewage treatment plants under aerobic conditions, the authors found EE2 to be largely persistent over an 80 hour treatment, while E2 and E1 were rapidly degraded. Moreover, in a natural situation (water from 3 rivers in England), the half-life of EE2 was calculated to be on average 17 days against 1.2 days for E2 (Jürgens et al., 2002). The stable concentrations of EE2 in our experimental setup also reveal that adsorption of EE2 to the polystyrene walls of the plate wells plays no role. Various types of materials and plastics can retain chemical compounds dissolved in water through adsorption (Walker & Watson, 2010). This can be a challenge in ecotoxicology studies if adsorption makes the tested compounds biologically unavailable and produces misleading results (Lung et al., 2000, Walker & Watson, 2010).

In our study, the high stability of EE2 concentrations in plates without embryos and its rapid decrease in plates with embryos indicates a quick and continuous uptake of the 2 pg EE2 by the embryo. When Bhandari et al. (2015) investigated EE2 uptake by embryonic stages of medaka, they also found significant uptakes. However, the authors exposed individuals to a much higher EE2 content of 50 pg per embryo, which is not likely in the wild (Bhandari et al., 2015). With an exposure content of 2 pg/embryo, estimated uptake in our experiment was continuous at around 0.5 pg per week and seemed to slow down after the first 4 weeks. This may be linked to the lower content of EE2 available in the water after 4 weeks of uptake. However, because of technical constraints (i.e. large water volumes are needed to determine low EE2 concentrations) we have only a single measurement per time point and treatment which limits our understanding on the measurement error. We can, nonetheless, estimate the measurement error based on the variance on measurements in plates without embryos (i.e. it was around 0.3 pg). The uptake of micropollutants may well be species dependent and vary with the chemical characteristics of the compounds. Lipophilic substances such as EE2 are taken up by eggs and embryos that are rich in lipids (Vermeirssen et al., unpublished data), such as salmonids eggs (Murzina et al., 2009).

A one-time exposure to 2 pg of EE2 was sufficient to reduce embryo survival by 0.7%.Given the high overall survival rate of nearly 98% in controls, this low reduction in survival represents a 35% increase in mortality. In our experimental setup embryos were singly raised under conditions that are arguably close to optimal for their development (e.g. minimizing pathogen growth and mechanical stress, etc.). In the wild, embryos are typically exposed to very strong selection (Elliott, 1994) and have to cope with a combination of challenges such as opportunistic microbes (Wilkins et al., 2015) and various types micropollutants (Chèvre, 2014, Moschet et al., 2014). If EE2-induced mortality is amplified by further stress factors (Wedekind et al., 2007, Segner et al., 2012, Segner et al., 2014), the results from our experimental treatments are likely to underestimate the potential ecotoxicological relevance of EE2.

The increased mortality we observed supports studies in typical model species (Caldwell et al., 2012, Aris et al., 2014) and the results of Brazzola et al. (2014), which found 1 ng/L of EE2 to reduce whitefish survival until hatching. In brown trout, Schubert et al. (2014) was unable to find a significant link between brown trout embryo mortality rates and E2 exposure. However, despite the high concentrations tested by the authors, E2 is naturally present in brown trout eggs and has a lower estrogenic potency than EE2 (Segner et al., 2003).

Embryos exposed to EE2 hatched slightly later (on average about 10 hours later) and at smaller size, i.e. EE2 reduced development and growth. Although small in an experimental setup, such differences in hatching time may create strong effects in the wild. Hatching time is fitness relevant because fast developing, and hence early emerging, salmonid embryos are more likely to establish and successfully defend a feeding territory than individuals that develop slower and emerge later (Einum & Fleming, 2000, Skoglund et al., 2012). Moreover, while embryos exposed to EE2 hatched at a smaller size, their yolk sac volume was not significantly larger than those of controls. This suggests that, more than only a delay in development, exposure to EE2 may have imposed an energetic cost on developing embryos. In accordance, EE2 has been described to change various aspects of development in typical model species (Caldwell et al., 2012, Aris et al., 2014) and to delay early development (Brazzola et al., 2014, Marques da Cunha et al., in prep.) and sexual differentiation (Selmoni et al., 2017) in salmonids.

EE2 is known to reduce growth of larvae of model species (Aris et al., 2014) and other salmonids. Brazzola et al. (2014) and Marques da Cunha et al. (in prep.), for example, have found a concentration of 1 ng/L EE2 to reduce larval growth in whitefish and European grayling (*Thymallus thymallus*), respectively. Here we found larval growth not to be significantly affected by the one-time addition of EE2 during early embryogenesis. These seemingly contrasting results may be caused by the different experimental protocols used, rather than revealing species-specific susceptibilities. While Brazzola et al. (2014) and Marques da Cunha et al. (in prep.) raised hatchlings under EE2 exposure until the last day of measurements, we transferred embryos to plates containing only standard water at the day of hatching. Moreover, from our chemical measurements, we conclude that there was nearly no EE2 left in the water at that time, i.e. our protocol did not allow for any further uptake of EE2 during the larval stages. The reduction of growth observed in Brazzola et al. (2014) and Marques da Cunha et al. (in prep.) may therefore be caused by an additional EE2 uptake by the larvae rather than revealing long-term effects of EE2 uptake during earlier embryonic stages.

In conclusion, our results show that embryos from a natural population of brown trout take up EE2 from water even at low and ecologically relevant concentrations. A total uptake of only 2 pg was sufficient to significantly affect embryo viability and development, despite the high levels of natural estrogens E1 and E2 in the embryos (estimated based on the increasing water content in Table S2). Ecotoxicology studies on EE2 and other micropollutants generally focus on concentrations that are kept constant during the assessments (Caldwell et al., 2012, Aris et al., 2014), i.e. they do not always consider exposure content and biological uptake. This can potentially lead to an underestimation of toxicities, as suggested by Quinnell et al. (2004). We found that an uptake of only 2 pg EE2 induces significant reduction in embryo viability in a species that is often exposed continuously to EE2 in their surrounding water. Our results suggest that toxicity of EE2 may be currently underestimated.

## Acknowledgements

We thank B. Bracher, U. Gutmann, and C. Küng, from the Fishery Inspectorate Bern for permissions and access to the fish, and L. Benaroyo, I. Castro, P. Christe, D. Maitre, D. Olbrich, C. Primmer, L. Wilkins, and D. Zeugin for assistance in the field/laboratory or discussions. This project was financially supported by Swiss National Science Foundation (grant to CW: 31003A_159579). This study complies with the relevant ethical regulations imposed by the University of Lausanne, the canton, and the country in which it was carried out.

## Author contributions

LMC, AU and CW designed the experiment on early trout viability, and LMC, AU, DN and CW performed the fieldwork. LMC and AU performed the experiment. ES and DN analyzed larval images. Data analysis was performed by LMC and CW. The experiment on EE2 measurements was designed by LMC, EV and CW. LMC, AU, DN and CW performed the fieldwork. The EE2 exposures and water collection was performed by LMC, and all chemical analyses were supervised by EV. LMC and CW wrote the first version of the manuscript that was then critically revised by all authors.

